# Metal Dependent Dynamic Equilibrium: A Regulatory Mechanism for M17 Aminopeptidases from *Plasmodium falciparum* and *Plasmodium vivax*

**DOI:** 10.1101/2020.10.08.331538

**Authors:** Tess R. Malcolm, Matthew J. Belousoff, Hariprasad Venugopal, Natalie A. Borg, Nyssa Drinkwater, Sarah C. Atkinson, Sheena McGowan

**Affiliations:** Infection & Immunity Program, Monash Biomedicine Discovery Institute and Department of Microbiology, Monash University, Clayton, Victoria, 3800, Australia; Ramacciotti Centre for Cryo-Electron Microscopy, Monash University, Clayton, Victoria, 3800, Australia; Infection & Immunity Program, Monash Biomedicine Discovery Institute and Department of Biochemistry and Molecular Biology, Monash University, Clayton, Victoria, 3800, Australia; Immunity and Immune Evasion Laboratory, Chronic Infectious and Inflammatory Diseases Research, School of Health and Biomedical Sciences, RMIT University, Bundoora, Victoria, 3083, Australia

**Keywords:** Leucine aminopeptidase, metalloprotease, oligomerization, regulation, *Plasmodium*, metallo-protease

## Abstract

M17 leucyl aminopeptidases are metal-dependent exopeptidases that rely on oligomerization to diversify their functional roles. The M17 aminopeptidases from *Plasmodium falciparum* (*Pf*A*-*M17) and *Plasmodium vivax* (*Pv*-M17) function as catalytically active hexamers to acquire free amino acids from human hemoglobin and are drug targets for the design of novel anti-malarial agents. In this study, we found that the active site metal ions essential for catalytic activity have a secondary structural role mediating the formation of active hexamers. We found that *Pf*A*-*M17 and *Pv-*M17 exist in a metal-dependent dynamic equilibrium between active hexameric species and smaller inactive species, that can be controlled by manipulating the identity and concentration of metal ions available. Mutation of residues involved in metal ion binding impaired catalytic activity and the formation of active hexamers. Structural resolution of the *Pv-*M17 hexameric species revealed that *Pf*A*-*M17 and *Pv-*M17 bind metal ions and substrates in a conserved fashion, although *Pv-*M17 forms the active hexamer more readily and processes substrates faster than *Pf*A*-*M17. On the basis of solution studies and structures determined by cryo-electron microscopy, we propose a dynamic equilibrium between monomer dimer tetramer hexamer, which becomes directional towards the large oligomeric states with the addition of metal ions. M17 aminopeptidases can exploit this sophisticated metal-dependent dynamic equilibrium to regulate formation of the catalytically active hexamer and therefore regulate catalysis.

## Introduction

The M17 family of metallo-aminopeptidases, often referred to leucine aminopeptidases (LAPs), selectively cleave N-terminal amino acids from polypeptide substrates (1). LAPs have varying roles in numerous essential processes across all kingdoms of life, including: regulation of immune responses in *Solanceae* (2,3), DNA recombination in *Escherichia coli* (4), free amino acid regulation in *Toxoplasma gondii* (5), and protection against oxidative stress in bovine cells (6). LAPs are routinely described as homo-hexamers, although variations in oligomeric states are observed in specific roles; monomers act as chaperones in tomatoes; catalytically active hexamers protect against oxidative stress; and multiple hexamers come together to aid in DNA recombination in *E. coli* (2,4,6,7).

LAPs have only ever been structurally resolved in the conserved homo-hexameric conformation. The catalytic C-terminal domains of each monomer are clustered in the center of the hexamer, to form a buried catalytic core, and are linked to the N-terminal domains by a central helical linker (8-11). Each active site has capacity to coordinate at least two metal ions, which are essential for proteolytic activity (12,13). The active site metal ions are coordinated at two distinct positions, denoted site 1 and site 2 (11,12). Although zinc ions are the most common occupants of both sites, site 1, or the loosely-bound site, can also coordinate Co^2+^, Mg^2+^, Mn^2+^, and Ca^2+^ (14). Site 2, or the tightly-bound site, is more limited and to date, has only been shown to only coordinate Zn^2+^ and Co^2+^ (12). Early studies suggested the two sites act independently to control enzyme function, with site 1 metal ions modulating rates of enzyme activity (*k*_cat_), and site 2 metal ions controlling enzyme activation through manipulation of substrate affinity (*K*_m_) (14,15). However, subsequent studies suggest the roles of site 1 and site 2 in enzymatic function may be more interdependent and complex than initially suggested (16). Despite their classification as zinc-metalloproteases, M17 aminopeptidases have been shown to have higher activity in the presence of cobalt or manganese ions than in the presence zinc ions (2,13,14,17,18).

The M17 aminopeptidases from the malaria-causing parasites, *Plasmodium falciparum* (*Pf*A-M17 or *Pf*LAP, (19) and *Plasmodium vivax* (*Pv*-M17 or *Pv*LAP, (20), have been functionally characterized and *Pf*A-M17 shown to be an attractive target for novel antimalarial drugs (21-25). In the *Plasmodium* parasite, M17 aminopeptidases are postulated to liberate free N-terminal amino acids from short hemoglobin-derived peptides for use in parasite protein production (21). *In vivo* localization studies indicate that *Pf*A-M17 and *Pv*-M17 function within the *Plasmodium* cytosol and *in vitro* analysis of aminopeptidase activity show that the two enzymes are most active in mildly basic solutions, reflective of the cytosolic environment (∼pH 8.0) (19,20). Both *Pf*A- and *Pv*-M17 cleave N-terminal leucine residues effectively (19,20), and the complete *Pf*A-M17 substrate selectivity profile indicates a preference for the hydrophobic amino acids leucine and tryptophan (26). Both aminopeptidases are present as hexamers in solution (19,20) and the *Pf*A-M17 crystal structure forms the conserved M17 family hexameric assembly (11).

The two active site metal ions within M17 aminopeptidases have previously been only described in terms of their necessity for catalytic function. In this study, we discovered that the active site metal ions of *Pf*A*-*M17 and *Pv-*M17 also play a structural role, operating as part of a previously undescribed mechanism of activity regulation for M17 aminopeptidases. We show that binding of active site metal ions mediates the association of inactive oligomers into active hexamers, and the dynamic equilibrium between those states could be manipulated with mutations designed to compromise active site metal binding. Structural characterization of the hexameric and tetrameric conformations using X-ray crystallography and cryo-electron microscopy, in combination with analytical ultracentrifugation (AUC) sedimentation velocity experiments, show that M17 aminopeptidases simultaneously adopt several oligomeric conformations and that the transition between oligomers (monomer dimer tetramer hexamer) is continuous, rapid and directionally controlled by the metal ion environment.

## Materials and Methods

### Molecular techniques

The gene encoding recombinant wild-type *Pv*-M17 (residues 73 – 621) with an in-frame C-terminal His^6^ tag was chemically synthesized by DNA 2.0 using codons optimized for gene expression in *E. coli. Pv-M17* was cloned into a pJ404 expression vector, which also encodes for ampicillin resistance. *Pv*-M17 mutants were generated by PCR mutagenesis. Metal binding mutant *Pv*M17-D323A was constructed and used as a template to produce double mutants, D323A E405L, utilizing primers outlined in Supplementary Table S1. The *Pv-*M17 N-terminal loop deletion mutant (*Pv*M17Δ53-78) was constructed using primer pairs amplifying outward from the region of interest (Supp. Table S1). The *Pf*A*-*M17 protein was produced from constructs described previously (11) and mutants *Pf*A-M17(AL) and *Pf*A*-*M17(W525A + Y533A) were synthesized by GenScript. Wild type and mutant sequences were confirmed using Sanger sequencing.

### Production and purification of Pv-M17 and PfA-M17

Recombinant *Pf*A-M17 and *Pv*-M17 genes were expressed in *Escherichia coli* BL21 DE3 cells. Cells were grown in auto-induction media to force overexpression of the target proteins. *E. coli* cells were lysed by sonication, centrifuged and the soluble supernatant was pooled. Proteins were purified via a two-step chromatography process involving nickel-affinity chromatography via the encoded C-terminal hexa-histidine tag (HisTrap™ Ni^2+^-NTA column; GE Healthcare Life Sciences), and size-exclusion chromatography (Superdex S200 10/300 GL column; GE Healthcare Life Sciences) as previously described (11,27). Proteins were stored in 50 mM HEPES pH 8.0, 0.3 M NaCl at −80°C until use.

### Enzyme kinetics

Aminopeptidase enzyme assays were carried out in white 384-well plates (Axygen) in a final volume of 50 µL. Aminopeptidase activity was determined by continually measuring the liberation of fluorogenic leaving group, 7-amido-4-methyl-coumarin (NHMec) from the commercially available substrate, Leucine-7-amido-4-methylcoumarin hydroxide (referred to in this study as Leu-Mec) (Sigma-Aldrich), using a FLUOStar Optima plate reader (BMG Labtech) with excitation and emission wavelength of 355 nm and 460 nm respectively. Emitted fluorescence was continuously recorded and quantified in fluorescence units. Data were analyzed using PRISM GraphPad 8 software. Metal ion and pH dependence were measured in 100 mM bis-Tris (pH 6.0, 7.0) or Tris (pH 8.0, 9.0), with 0.2 mM or 1.0 mM MnCl_2_, MgCl_2_, ZnCl_2_ and CoCl_2_. 150 nM *Pf*A-M17 or *Pv-*M17 was incubated with metal ions for 10 minutes at 37°C before addition of 10 µM substrate and activity measured for one hour, or until steady-state was achieved. Kinetic assays were carried out in 100 mM Tris pH 8.0 buffer supplemented with 1 mM CoCl_2_. *Pf*A-M17 and *Pv*-M17 concentrations were constant at 150 nM. Substrate concentration ranged from 0 µM to 500 µM. Substrate was incubated for 10 mins at 37°C before addition of protein and activity measured for one hour.

Hydrolysis of the cysteinyl-glycine dipeptide substrate (referred to as Cys-Gly in this study) was measured using a colorimetric method detecting liberation of free cysteine from the Cys-Gly dipeptide, using a protocol adapted from (6). Reactions were carried out in 1.5 mL microcentrifuge tubes and reactions volumes were 100 µL. 250 nM of *Pf*A-M17/*Pv*-M17 was incubated with 2 mM Cys-Gly at 37°C for 15 minutes in varying metal ions at 0.2 mM (Zn, Mn, Co, Ni, Mg, Metal free) and pH conditions (bis-Tris pH 6.0, bis-Tris pH 7.0, Tris-Cl pH 8.0, Tris-Cl pH 9.0). Reactions were stopped by the addition of 100 µL 100 % acetic acid. 100 µL of a ninhydrin reagent solution containing 50 mg ninhydrin dissolved in 2 mL 4 M HCl/acetic acid (2:3) was added and tubes were lightly mixed and centrifuged using a benchtop microcentrifuge. Samples were heated at 100°C for 10 minutes, then cooled on ice before 100 µL was transferred to a 96-well plate and absorbance measured at 560 nm (Clariostar, BMG LabTech).

### Analytical Size-Exclusion Chromatography

A 100 µL sample was loaded on to a Superdex S200 Increase 10/300 GL gel-filtration column (GE Healthcare Life Sciences) pre-equilibrated with 50 mM HEPES pH 8.0, 0.3 M NaCl using a Fast Protein Liquid Chromatography (FPLC) system (ÄKTA Purifier, GE Healthcare Life Sciences). Prior to loading, concentrated samples were centrifuged at 13,000 RPM, 20 mins, 4°C to remove any residual debris or aggregate. A flow-rate of 0.5 mL/min was applied for a minimum of 25 mL and 0.5 mL fractions collected (Äkta Frac-920, GE Healthcare Life Sciences). Size Exclusion Chromatography Multi-Angle Light Scattering was carried out in identical conditions to analytical SEC experiments (UFLC, Shimadzu; Dawn Helios-II, Wyatt; Optilab T-rEX, Wyatt) and data was analyzed using Astra 6. To test fraction activity, 25 µL of each fraction (concentration not normalized) was pipetted into a white 384-well plate and incubated at 37°C for 10 mins. Following incubation, 10 µM Leu-Mec substrate diluted in 100 mM Tris pH 8.0, 1.0 mM CoCl_2_ was added and activity monitored for 60 mins. Resultant activity across the 60-minute time frame was expressed as a fluorescence units/min and plotted against corresponding fraction elution volume using Prism GraphPad 8.

### Analytical Ultra Centrifugation

Analytical ultracentrifugation (AUC) experiments were conducted using a Beckman Coulter Optima Analytical Ultracentrifuge at a temperature of 20 °C. For sedimentation velocity experiments, samples were loaded at a concentration of 0.5 mg/mL. Sample buffer (50 mM HEPES pH 8.0, 0.3 M NaCl) was supplemented with 50 µM Mn^2+^, Co^2+^, Mg^2+^, Zn^2+^ or 1 mM Mn^2+^ as appropriate. 380 μL of sample and 400 μL of reference solution (sample buffer) were loaded into a conventional double sector quartz cell and mounted in a Beckman 4-hole An-60 or 8-hole An-50 Ti rotor. Samples were centrifuged at a rotor speed of 40,000 rpm and the data was collected continuously at a single wavelength, most commonly at 280 nm. Solvent density and viscosity, as well as estimates of the partial specific volume (0.742928 mL/g for *Pf*A-M17, 0.739998 mL/g for *Pv*-M17 at 20 °C) were computed using the program SEDNTERP (28). Sedimentation velocity data were fitted to continuous size [*c*(*s*)] distribution model using the program SEDFIT (29,30). Data was plotted using Prism Graphpad 8.

For sedimentation equilibrium experiments, reference and sample sectors were loaded with 140 μL reference and 100 μL sample (8.3 µM and 4.2 µM) plus 20 μL FC-43 oil in the sample sector. After initial scans at 3,000 rpm to determine optimum wavelength and radial range for the experiment, samples were centrifuged at 12,000 and 18,000 rpm at 20 °C. Data at each speed were collected at multiple wavelengths every hour until sedimentation equilibrium was attained (24 h). Sedimentation equilibrium data were analyzed with SEDPHAT (31) using the single species analysis model.

### X-ray crystallography of Pv-M17 N-terminal mutant

N-terminal loop deletion *Pv*M17Δ53-78 crystallization conditions were identified using sparse-matrix crystal screening. Crystals were observed in 0.1 M HEPES pH 6.5, 15% PEG 3350 and 0.15 M (NH_4_)_2_SO_4_ at protein concentrations of 3 and 6 mg/mL, and reservoir: protein ratio of 1:1. Crystals were mostly singular and rectangular prism in morphology. Data were collected at 100 K at the Australian Synchrotron MX2 beamline 31D1 (32). Diffraction data was collected to 2.6 Å. Diffraction images were processed using X-ray detector software (XDS) and Pointless (33,34). 5% of the reflections were set aside for calculation of R_*free*_ to compare against refined data. The *Pv-*M17Δ53-78 structure was solved by molecular replacement using a search model constructed from the *Pf*A-M17 crystal structure (RCSB ID 3KQZ) in Chainsaw (35). A molecular replacement search using Phaser (36,37) identified twelve copies of the search model in the asymmetric unit, arranged into two independent hexamers. Model building proceeded from the initial Phaser output using PHENIX refinement (38). For initial refinement rounds in PHENIX, non-crystallographic symmetry (NCS) maps were produced to assist in initial model building. Between rounds of refinement, the structure was visualized using COOT (39,40), and the structure adjusted according to 2*F*o-*F*c and *F*o-*F*c electron density maps. After several iterations of refinement and model building, NCS maps were disabled as an option in PHENIX, and the stereochemistry and ADP weight optimized. On occasion, an NCS map was calculated in COOT to assist in modelling of flexible regions and placement of ligands. The structure was validated with Molprobity (41) and figures generated using PyMOL version 1.8.2.3.

### Cryo-electron microscopy of Pv-M17 oligomeric states

For the single particle cryo-EM analysis of the hexameric and smaller oligomeric conformations, *Pv-*M17 was concentrated to ∼ 2 mg/mL and pre-incubated in the presence of 1 mM MnCl_2_ or 100 mM EDTA respectively. 4 µL of sample was applied to Quantifoil Cu R 1.2/1.3 grids (Quantifoil Microtools GmbH) which has undergone a plasma discharge and were vitrified in liquid ethane using a Vitrobot MkIV plunge freezer (Thermo Fisher Scientific), with the sample chamber set to 100% humidity and 4°C. Grids were stored in liquid nitrogen until data collection. Data were collected on a FEI Titan Krios (Thermo Fisher Scientific) operating at 300 kV fitted with a Quantum energy filter (Gatan) and K2 (Gatan) Direct Electron Detector. Data was collecting using a pixel size of 1.06 Å and with a total average electron dose per image of 63 e/Å^2^ for the hexameric species and 105 e/Å^2^ for the tetrameric species.

Micrograph movies were imported into RELION (v3.0, v3.1) (42) and motion corrected using MotionCor2 and CTF estimation performed using CTFFIND4.1 (43). Particles were picked using the auto-picking function in RELION using a Laplacian-of-Gaussian approach. Extracted particles were analyzed using RELION and cryoSPARC (44), and reference free 2D classification was performed. 2D Classes that corresponded to the expected *Pv-*M17 molecule were selected and an ab initio 3D model was generated in CryoSPARC. This model was used as an initial model for further Euler angler refinement in RELION.

Final 3D maps were further refined using RELION using the Gold-standard approach to a final resolution of 2.8 Å. Maps were inspected using UCSF Chimera (45). For the hexameric structure, the *Pv*M17Δ53-78 crystal structure determined as part of this study was used as an atomic model template and fitted to the cryo-EM electron density map. Adjustments to the template model were carried out using Coot and the final hexamer model refined using the real-space refinement tool in PHENIX. A tetramer based on the hexamer model was used as the starting model for the low oligomeric weight map. The starting model was docked into target map using Chimera ‘Fit in Map’ function. Molecular dynamic flexible fitting (MDFF) simulations was used to produce a reasonable starting atomic model and was performed using NAMD (v2.13) software package (46). Final model refinements were performed by real space refinement as implemented in the PHENIX software package (38) followed by manual model curation in COOT (39,40). See Supplementary Table S2 for structure statistics.

## Results

### PfA-M17 and Pv-M17 exhibit unique oligomerization behaviors

Recombinant *Pf*A-M17 and *Pv*-M17 were successfully purified from a bacterial expression system using a two-step chromatography method. The purified proteins were active in previously reported assay conditions (19,20) and had similar kinetic parameters to those previously reported (Table 1). Interestingly, the *Pv*-M17 enzyme is significantly faster than *Pf*A-M17 with *k*_cat_ values at least 10x higher than *Pf*A-M17 (Table 1). During purification, both *Pv-*M17 and *Pf*A-M17 eluted from SEC as broad peaks suggestive of poorly separated oligomeric species. To further investigate the solution species of each protein, SEC-MALS experiments with 100 µL samples at 3 mg/mL were used to dissect the oligomeric content of the protein samples. At the lower concentration and volume, *Pf*A-M17 separated into two distinct populations at molar masses of ∼380 kDa and ∼70 kDa, which approximately correlated to a hexamer and monomer respectively (Fig 1A). The peak area shows that *Pf*A-M17 predominantly adopted the monomeric conformation with only a small proportion forming the hexamer. *Pv-*M17 also separated into two distinct populations, at a high molar mass (∼330 - 370 kDa) spanning the hexamer molecular weight and at a low molar mass (∼120 kDa), correlating to a dimer (Fig 1B). These two populations were of approximately equal concentration and connected by a non-zero baseline, indicating dynamic movement between the two populations (Fig 1B, gray bars). Catalytic activity of the separated *Pf*A-and *Pv*-M17 oligomeric states was almost completely limited to the hexameric populations (Fig 1A & B). The small oligomeric species display some residual activity, however, this may be due to the small oligomers forming larger active species throughout the course of the experiment. More distinct separation of the *Pv*-M17 and *Pf*A-M17 oligomeric states was achieved using AUC sedimentation velocity experiments. Higher resolution separation of *Pv-*M17 showed at least three distinct populations at standardized (*s*_20,w_) sedimentation coefficients of ∼6S, ∼9S and ∼12S, with dynamic movement between these three populations (Fig 1C). AUC analysis of *Pf*A-M17 suggested the small molar mass peak observed in SEC-MALS experiments actually consists of two distinct populations at *s*_20,w_ of approximately 4S and 6S (Fig 1C).

**Table 1.**
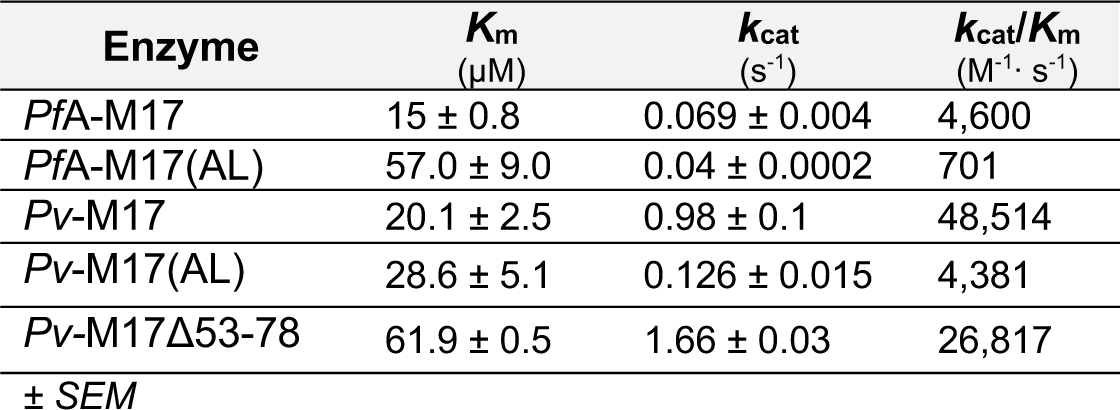
Kinetic characterization of *Pf*A*-*M17 and *Pv-*M17 wild types and mutants against fluorescent substrate, Leu-Mec.

**Figure 1.**
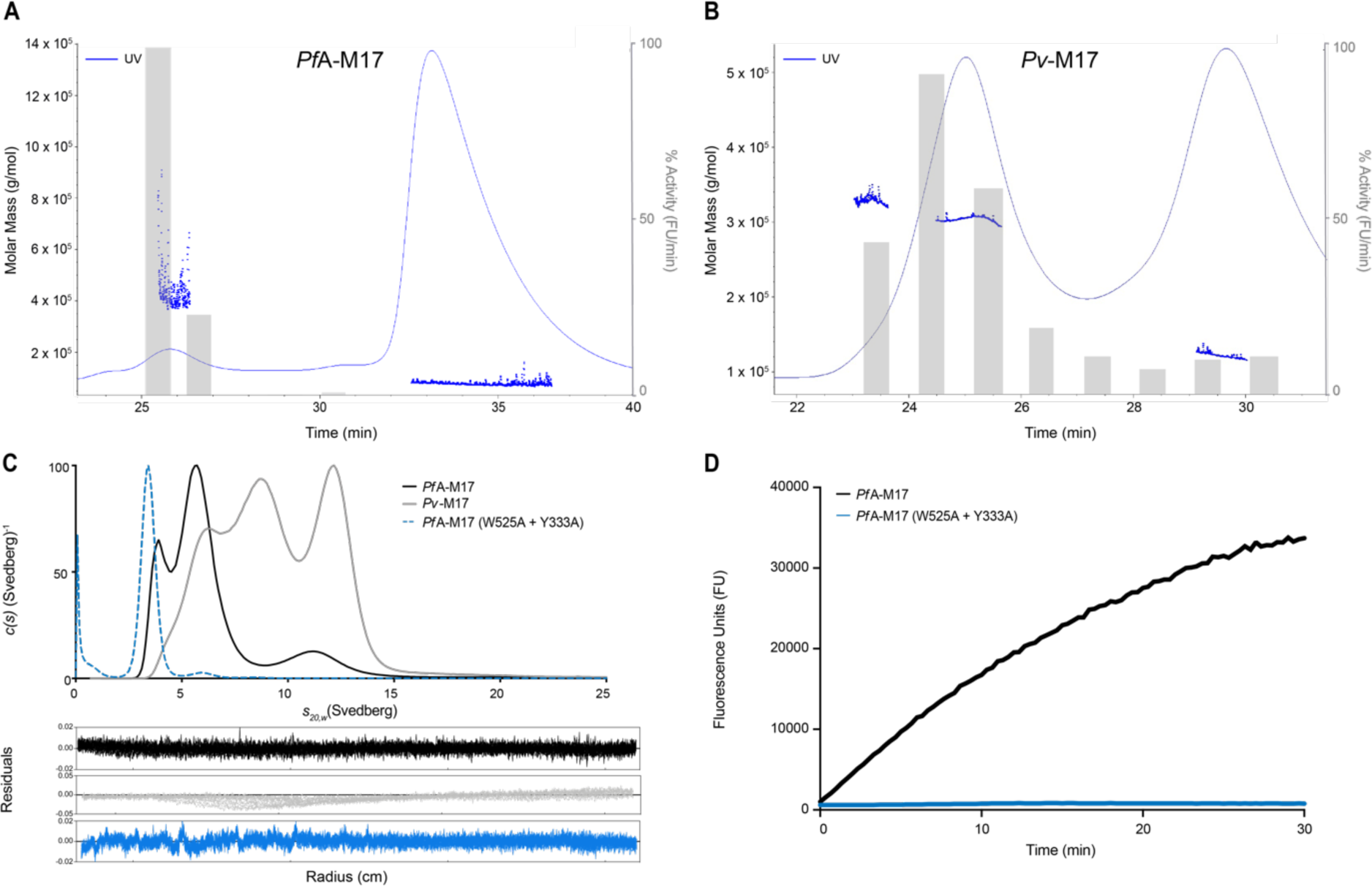
*Pf*A-M17 and *Pv*-M17 exist as a mix of oligomeric states in solution. FPLC SEC-MALS traces of **(A)** *Pf*A-M17 and **(B)** *Pv*-M17 in 50 mM HEPES pH 8.0, 0.3 M NaCl with approximate sizes calculated using SEC-MALS. Activity of each fraction was measured as an activity rate (fluorescence units/min), normalized and represented by grey bars. Both *Pf*A-M17 and *Pv*-M17 are present as a mix of oligomeric states in solution, however only the large oligomeric state shows substantial catalytic activity. **(C)** *c*(*s*) distribution analysis of sedimentation velocity experiments of *Pf*A-M17 (black), *Pv*-M17 (grey) and *Pf*A*-*M17 (W525A + Y533A) in 50 mM HEPES pH 8.0, 0.3 M NaCl. Both *Pf*A-M17 and *Pv*-M17 sedimented as multiple species, whereas *Pf*A*-*M17 (W525A + Y533A) sedimented as a single monomeric species. Distributions were normalized so that all had a maximum of 100. **(D)** Comparative activity levels of *Pf*A*-*M17 wild type (black) and *Pf*A*-*M17 (W525A + Y533A) (blue). Enzyme concentrations were uniform between the two samples, and activity levels measured using fluorescence units (FU). *Pf*A*-*M17 wild type is active, whereas the *Pf*A*-*M17 (W525A + Y533A) is inactive.

To create homogenous monomeric or stable small oligomeric species for use as a reference in AUC experiments, we sort to introduce point mutations to disrupt hexamerization. Investigation of our *Pf*A*-*M17 structure identified that Trp525 and Phe533 formed two sets of pi-pi stacking interactions with corresponding residues from another chain within the hexamer. We hypothesized that these two residues were likely involved in stabilizing the hexameric conformation. We mutated both residues to Ala to eliminate these aromatic interactions between subunits. In solution, the *Pf*A*-*M17 (W525A + Y533A) protein formed a single species that had a sedimentation coefficient of ∼3.7S (Fig 1C), which we confirmed as monomeric using sedimentation equilibrium experiments (Supp. Fig S1). Monomeric *Pf*A*-*M17 (W525A + Y533A) was enzymatically inactive compared to wild type *Pf*A-M17 in the same assay conditions (Fig 1D). From this result, we could predict that the *Pf*A*-*M17 WT sample in Fig 1C contained monomeric species at 4S and dimeric species at 6S, while *Pv-*M17 contained the dimeric species at 6S, and tetrameric and hexameric species at 9S and 12S respectively.

### Oligomerization and activity are mediated by active site metal ions

With a relationship between the hexameric conformation and catalytic activity confirmed, we were interested in investigating factors that mediate the association of small inactive oligomers into active hexamers. M17 aminopeptidases are metal-dependent proteases that coordinate at least two metal ions per active site (11,12). Our analysis of purified *Pf*A-M17 showed that it was predominantly monomeric in solution when purified from a bacterial expression system and that this monomer was catalytically inactive (Fig 1A, 1D). However, previous studies by us (11) and others have shown that addition of a divalent cation to assay buffers converts the inactive species to an active aminopeptidase. Therefore, we postulated that metal ions may also be playing an essential role in the formation of active hexamers. To investigate the influence of different metal ions on oligomerization and activity, both AUC sedimentation velocity experiments and catalytic activity assays were conducted in the presence of biologically relevant metal ions (Zn^2+^, Mg^2+^, Mn^2+^ and Co^2+^). We observed unique oligomerization and activity responses to each metal ion tested, and an overall behavioral difference between *Pv*-M17 and *Pf*A-M17. *Pf*A-M17 did not completely hexamerize in any of the metal ion conditions tested (Fig 2A), although Mn^2+^ and Co^2+^ did result in a shift toward the larger oligomeric states. Mg^2+^ failed to induce any oligomerization of *Pf*A*-*M17 given the AUC sedimentation profile remained similar to that of the no metal added sample. *Pv-*M17 associated into a single species in the presence of Mn^2+^ and Co^2+^ with sedimentation coefficients at 12S and 13.5S respectively (Fig 2B). Mg^2+^ caused a more defined separation of different oligomeric species although had minimal effect on overall oligomeric species present. Using AUC sedimentation equilibrium experiments, the Mn^2+^ -induced 12S species was confirmed to have a molar mass of ∼356 kDa, consistent with the active hexamer (Supp. Fig S2, Supp. Table S3). Zn^2+^ was also tested as part of these experiments for both proteins, however the proteins precipitated at low metal ion concentration.

**Figure 2.**
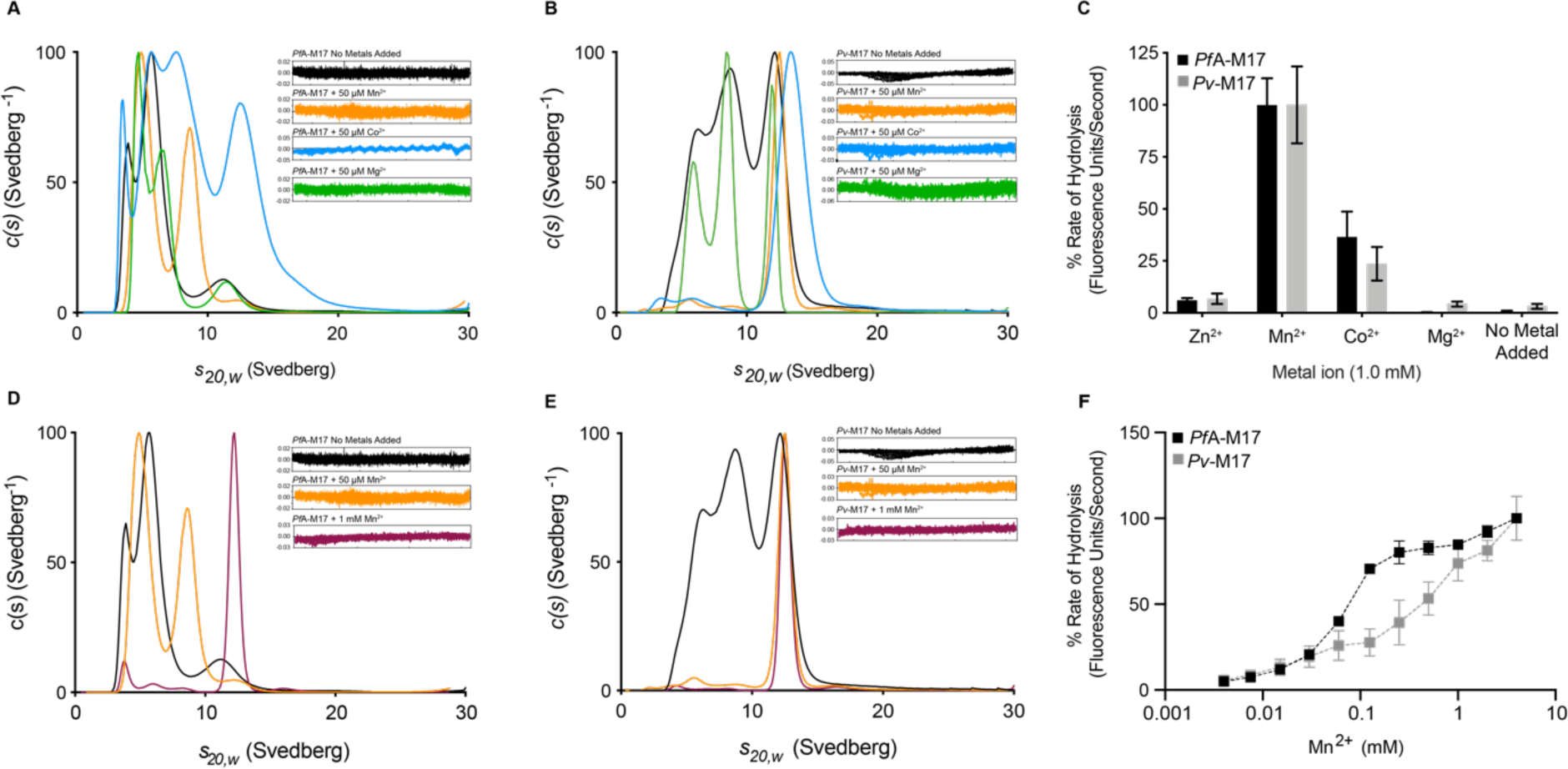
Catalytic activity and oligomerization are influenced by environmental metal ion identity and concentration. **(A**) *c*(*s*) distribution analysis of sedimentation velocity experiments of *Pf*A-M17 in the presence of no added metal (black), 50 µM Mn^2+^ (orange), 50 µM Co^2+^ (blue) or 50 µM Mg^2+^ (green). *c*(*s*) values are normalized with the highest *c*(*s*) value represented by 100%. Residuals for each experiment in inset. Mn^2+^ and Co^2+^ cause *Pf*A*-*M17 to shift towards large oligomeric states. **(B)** *c*(*s*) distribution analysis of sedimentation velocity experiments of *Pv*-M17 in the same conditions as (A). *c*(*s*) values are normalized with the highest c(s) value represented by 100%. Residuals for each experiment in inset. Mn^2+^ and Co^2+^ cause *Pv-*M17 to hexamerize. **(C)** Activity of *Pf*A-M17 (black) and *Pv-*M17 (grey) in presence of 1.0 mM divalent metal ions and no metal ions added. Activity rates are normalized to highest activity rate for each enzyme. Activity rates are highest for both *Pf*A*-*M17 and *Pv-*M17 in the presence of Mn^2+^ and Co^2+^. **(D)** *c*(*s*) distribution analysis of sedimentation velocity experiments of *Pf*A-M17 in the presence of no metal added (black), 50 µM Mn^2+^ or 1 mM Mn^2+^ (maroon). *c*(*s*) values are normalized with the highest *c*(*s*) value represented by 100%. Residuals for each experiment in inset. *Pf*A-M17 reaches full hexamerization in the presence of 1 mM Mn^2+^ **(E)** *c*(*s*) distribution analysis of sedimentation velocity experiments of *Pv*-M17 in the same conditions as D. *c*(*s*) values are normalized with the highest *c*(*s*) value represented by 100%. Residuals for each experiment in inset. *Pv-*M17 hexamerizes in Mn^2+^ concentrations as low as 50 µM. **(F)** *Pf*A-M17 (black) and *Pv-*M17 (grey) catalytic activity in increasing Mn^2+^ concentrations. Activity of both aminopeptidases increases as metal ion concentration increases.

Catalytic activity was also tested using the same panel of biologically relevant metal ions. Both *Pf*A-M17 and *Pv-*M17 exhibited the highest activity in the presence of Mn^2+^ and Co^2+^, and negligible activity in Mg^2+^ and Zn^2+^ (Fig 2C). When compared with AUC data, the metal ions that induced a complete or partial oligomeric shift towards the hexameric conformation were the same ions that corresponded with high catalytic activity levels. Conversely, buffer conditions that produced relatively small proportions of the hexameric species, resulted in comparatively low catalytic activity levels.

Given both *Pf*A*-*M17 and *Pv-*M17 demonstrated a shift towards higher order oligomers coupled with high activity levels in the presence of Mn^2+^, we were interested to see if metal ion concentration, in conjunction with metal ion identity, contribute to formation of the active hexameric species. AUC sedimentation velocity experiments were carried out in increasing Mn^2+^ concentrations; 0 µM, 50 µM and 1 mM. As metal ion concentration increased we observed an increase in the concentration of active hexamer present in solution. *Pf*A*-*M17 exhibited a slight shift towards the larger oligomeric states in the presence of 50 µM and required 1 mM Mn^2+^ to achieve complete hexamerization. (Fig 2D) Conversely, *Pv-*M17 hexamerized readily in 50 µM Mn^2+^ and became a more defined peak in the presence of 1 mM Mn^2+^ (Fig 2E). We also assessed catalytic activity of both *Pf*A-M17 and *Pv-*M17 in the presence of increasing Mn^2+^ concentration (0.004 mM – 4.0 mM). Activity rates positively correlated with the increasing Mn^2+^ concentration for both *Pf*A*-*M17 and *Pv-*M17, supporting the AUC results to show that increasing concentrations of metal ions increases the concentration of catalytically active hexamers. Again, *Pf*A*-*M17 shows a distinct oligomerization/activity profile compared to *Pv-*M17. *Pf*A*-*M17 activity appears to plateau at [Mn^2+^] = 0.5 mM, before continuing in an upward trajectory, whereas *Pv-*M17 reaches the first activity plateau at [Mn^2+^] = 0.1 mM and actually reaches a second plateau at [Mn^2+^] = 1 mM before again rising in activity. Maric et al. (17) described a similar activity profile in increasing Zn^2+^ concentrations and suggested the plateaus were due to saturation of the “loosely-bound” and “tightly-bound” metal-binding sites. By this model, *Pv-*M17 reached saturation of the loosely bound site at lower metal ion concentrations than *Pf*A*-*M17, suggesting that *Pv-*M17 may coordinate metal ions in this position more readily.

### Active site metal ion binding is essential for oligomerization

To confirm that the correlation between metal ion environment/concentration and hexamer formation was due to metal binding within the active site of the enzymes, we introduced point mutations designed to impair the ability of *Pf*A*-*M17 and *Pv-*M17 to bind active site metal ions. We chose to alter the conserved Asp379 and Glu461 residues that show bidentate coordination of both metal ions to prevent or hinder the active site from functioning as a metal accepting pocket. These residues were changed to Ala and Leu respectively, producing mutants *Pf*A*-*M17 (D379A + E461L) and *Pv-*M17 (D323A and E405L), which we refer to herein as *Pf*A-M17(AL) and *Pv*-M17(AL) respectively. The mutants were purified by the standard protocol and exhibited no obvious differences to wild-type enzymes(s) during purification. The catalytic efficiency of both *Pf*A-M17(AL) and *Pv*-M17(AL) was approximately 10-fold lower than the respective wild type enzyme (Table 1). In *Pv-*M17, this can be attributed primarily to the marked reduction in substrate turnover (*k*_cat_) as opposed to substrate binding (*K*_m_), which was largely unchanged compared to wild type. However, *Pf*A-M17 substrate affinity was lower in *Pf*A-M17(AL) than wild-type, which likely contributes to decreased turnover rates and overall reduced catalytic efficiency (Table 1).

Oligomerization of *Pf*A-M17(AL) and *Pv*-M17(AL), compared to WT, was tested using AUC sedimentation velocity experiments in the presence and absence of 1 mM Mn^2+^. In the absence of Mn^2+^, *Pf*A-M17(AL) sedimented as a very broad peak at 10S with a diminishing tail stretching to ∼ 4S and a second, much smaller peak at ∼4.5S (Fig 3A), suggesting that the aminopeptidase was in a state of rapid and dynamic transition between low and high order oligomers. When 1 mM Mn^2+^ was added, the *Pf*A*-*M17(AL) *c*(*s*) distribution became broader and absorbed the smaller peak observed earlier at ∼4.5S but remained largely unchanged with the addition of metal ions. (Fig 3A). *Pv-*M17(AL) showed similar oligomerization characteristics to *Pf*A*-*M17(AL), adopting a broad curve that peaked at ∼11S with a diminishing tail that extended to ∼4S (Fig 3B). Addition of 1 mM Mn^2+^ caused the *Pv-*M17(AL) *c*(*s*) distribution to broaden slightly, but as seen with *Pf*A*-*M17(AL), the distribution remained largely unchanged (Fig 3B). Distinct hexamerization is observed with addition of Mn^2+^ to *Pf*A*-*M17 WT and *Pv-*M17 WT (Fig 3A and 3B). In contrast, *Pf*A*-*M17(AL) and *Pv-*M17(AL) both showed broad sedimentation profiles regardless of the presence of metal ions, suggesting that *Pf*A*-* M17 (AL) and *Pv-*M17 (AL) fluctuate across a broad range of oligomeric species when metal binding is impeded. Furthermore, *Pf*A*-*M17(AL) and *Pv-*M17(AL) adopted very similar oligomerization profiles when metal binding was impeded, unlike *Pf*A*-*M17 WT and *Pv-*M17 WT which demonstrate distinct profiles when compared.

**Figure 3.**
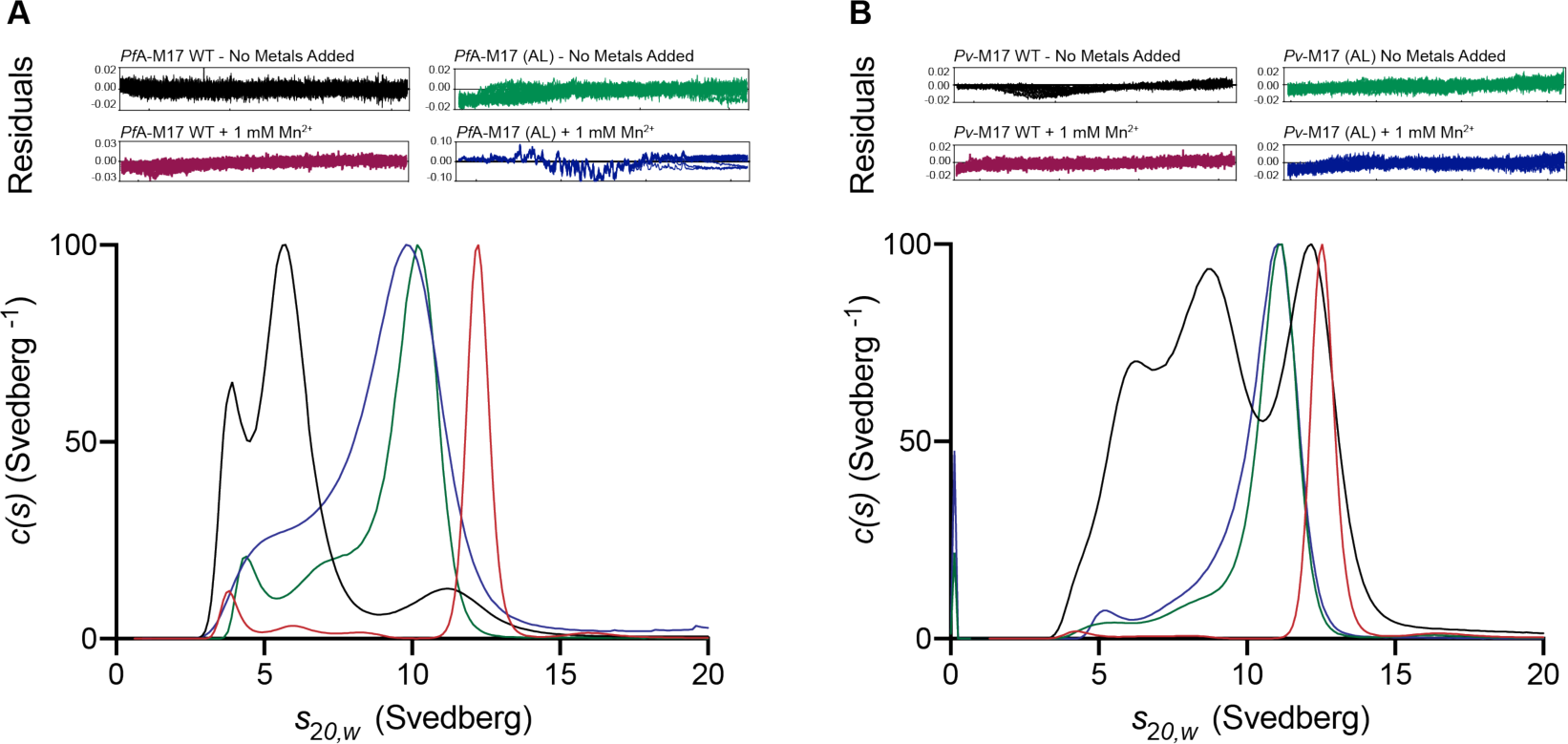
Compromising active site metal binding disrupts the oligomerization mechanism and reduces catalytic activity. *c*(*s*) distribution analysis of sedimentation velocity experiments of *Pf*A-M17 (A) and *Pv*-M17 (B). Wild type M17 in absence of Mn^2+^ (grey) and presence of Mn^2+^ (black). *Pf*A*-*M17 (AL) and *Pv-*M17 (AL) in absence of Mn^2+^ (red) and presence of Mn^2+^ (blue).

### Cysteinyl-glycinase activity is mediated by a two-fold metal-dependent mechanism

M17 family aminopeptidases have been shown to digest the Cys-Gly dipeptide as part of the glutathione regulation pathway but only in the presence of manganese (6,7,47). We therefore investigated in addition to influencing oligomerization. We characterized the cysteinyl-glycinase activity of *Pf*A-M17 and *Pv-*M17 against the Cys-Gly dipeptide at pH ranging from 6.0 to 9.0, and in the presence of Zn^2+^, Mn^2+^, Co^2+^, Mg^2+^ and also with no metal ions added. *Pf*A*-*M17 only showed catalysis of Cys-Gly in the presence of Mn^2+^ at a pH level of 8.0 (Fig 4A). *Pv-*M17 also demonstrated activity in Mn^2+^ at pH 7.0 and 8.0 (Fig 4B). *Pv-*M17 maintains a faster rate of catalysis than *Pf*A*-*M17; *Pv-*M17 reached absorbance levels in excess of 1.0 mAu while *Pf*A*-*M17 reached absorbance levels of just 0.2 – 0.3 mAu in the same time frame. While *Pf*A*-*M17 and *Pv-*M17 were both active against Leu-Mec in the presence of Co^2+^ at pH 8.0 (Fig 2C), neither exhibited catalysis of Cys-Gly in the same conditions despite the presence of the active hexameric conformation (Fig 4A, B). Despite the change in substrate from earlier characterization experiments, *Pv-*M17 maintained a faster catalysis rate than *Pf*A*-* M17 (Fig 4A, B).

**Figure 4.**
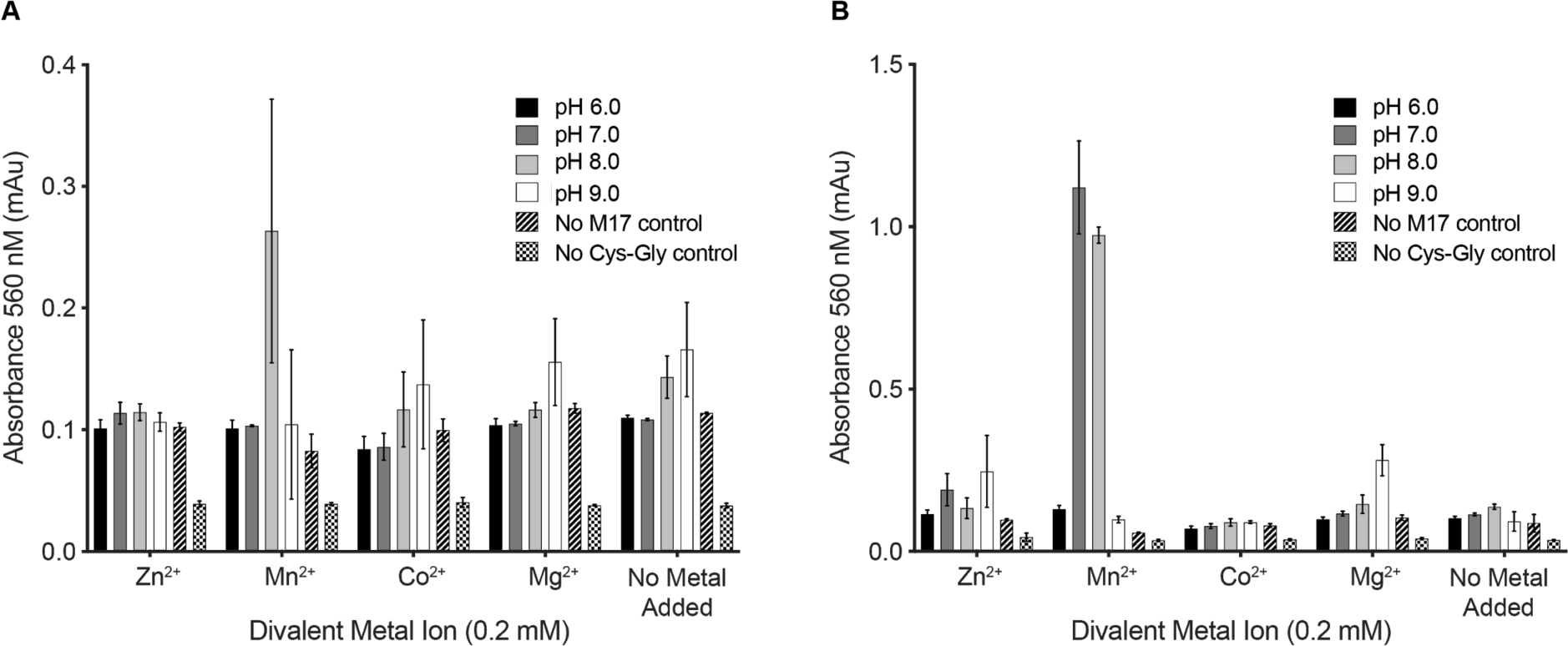
*Pf*A*-*M17 and *Pv-*M17 hydrolyze the cysteinyl-glycine dipeptide. Activity of **(A)** *Pf*A*-*M17 and **(B)** *Pv-*M17 against cysteinyl-glycine dipeptide in the presence of 0.2 mM Zn^2+^, Mn^2+^, Co^2+^, Mg^2+^ and with no metal ions added in buffer varying pH from 6.0 to 7.0. Amount of liberated cysteine was detected using ninhydrin and the resultant absorbance measured at 560 nm. Both *Pf*A*-*M17 and *Pv-*M17 are only active against the dipeptide in the presence of Mn^2+^, *Pf*A*-*M17 at pH 8.0 and *Pv-*M17 at pH 7.0 and pH 8.0.

### PfA-M17 and Pv-M17 unique behavior is not due to differences in active site structure

Having continually observed differences in the metal-responsive behavior between *Pf*A-M17 and *Pv-*M17, we were interested to investigate whether there were any structural differences in the active sites that would lead to disparities in metal binding and therefore in oligomerization and activity. The X-ray crystal structure of *Pf*A-M17 was solved prior to this study, with all structures – both unliganded unbound and in complex with various inhibitors (11,24,48,49) - showing the hexameric conformation that is characteristic of M17 family aminopeptidases. After failing to obtain high-resolution X-ray data of full length *Pv-*M17, we identified a unique 21 amino acid insertion rich in Gly and Ser (Gly = 42.8%, Ser = 33%) within the postulated N-terminal region (Supp. Fig S3). The insertion which we have called the ‘*Pv* loop’ was predicted to be a solvent-exposed flexible loop that may impede crystal packing and diminish data quality. To overcome this, the 25 amino acid *Pv* loop was excised from the N-terminal domain to produce the mutant *Pv*-M17Δ53-78.

*Pv*-M17Δ53-78 was successfully crystallized and a high-resolution X-ray crystal structure solved to 2.6 Å (PDB ID: 6WVV) (Supp. Table S4). The overall structure adopted the characteristic M17 hexameric conformation, composed of a ‘dimer of trimers’ (Fig 5A) with the six chains clustered within the hexamer core, with each catalytic domain connected to a solvent-exposed N-terminal domain (Fig 5A). There were two complete hexamers in the asymmetric unit, with an overall RMSD over 2256 Cα atoms of 0.194 Å. Minor rigid body movement of the N-terminal domains relative to each other was evident upon overlay of each chain (Supp. Fig S4A). Several solvent-exposed regions (Ser197 – Glu199, Ala305 – Glu309 in all chains, and Glu98 – Asn100 and Lys122 – Val124 in chains G-L) could not be modelled, suggesting these regions are disordered.

**Figure 5.**
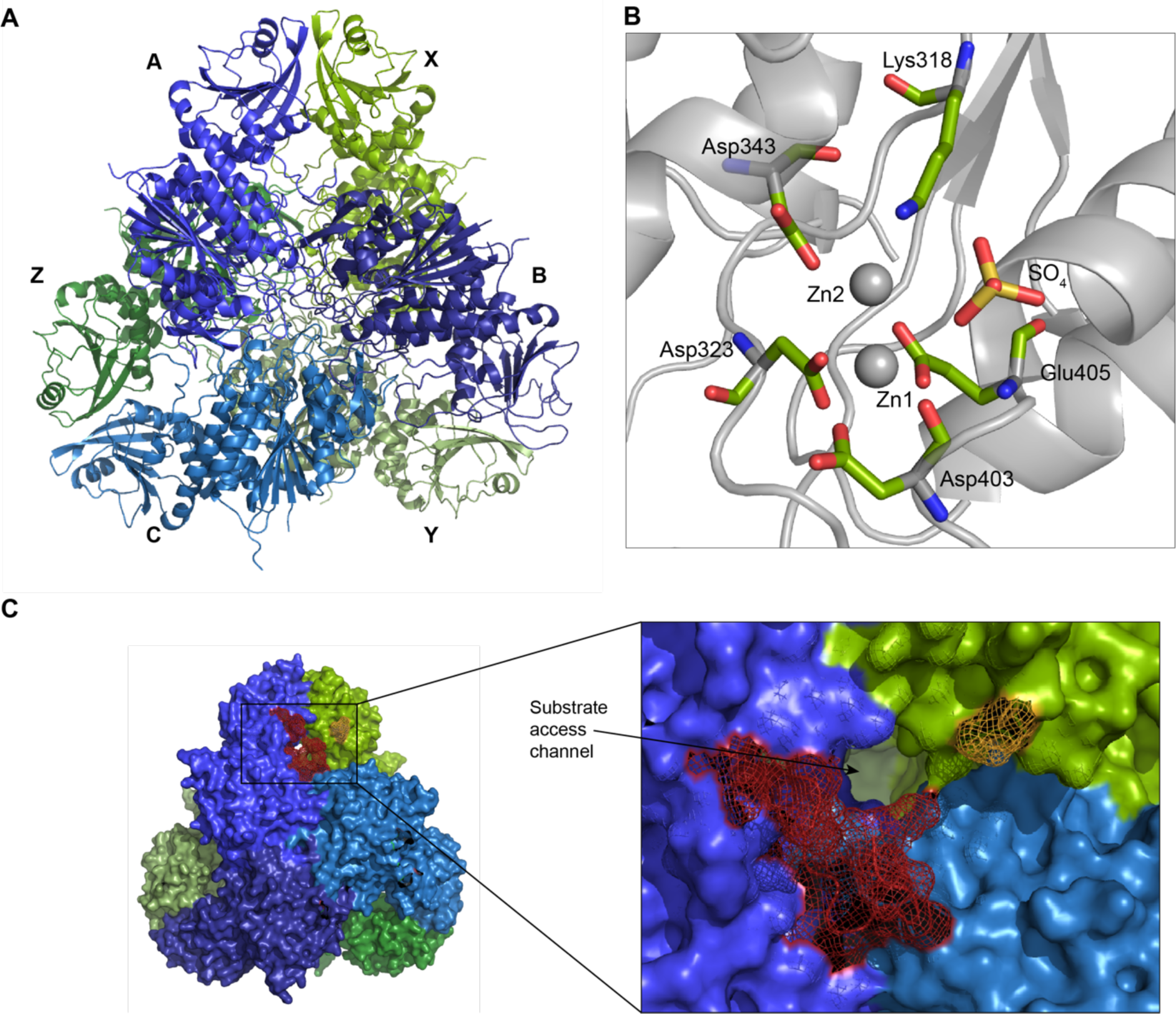
The *Pv-*M17Δ53-78 X-ray crystal structure adopts the conserved hexameric conformation. **(A)** 2.6 Å X-ray crystal structure of *Pv-*M17Δ53-78 adopts a hexameric conformation, composed of a ‘dimer or trimers’. Front trimer is depicted in shades of blue and chains are labeled A, B and C. Back trimer is depicted in shades of green and chain are labeled X, Y and Z. **(B)** *Pv-*M17Δ53-78 X-ray crystal structure active site. Metal-coordinating residues are shown as green sticks, colored by atom. Zn1 and Zn2 are depicted as spheres. Zn1 is coordinated by the carboxyl group of Asp323, Asp403 and Glu405 and the amine group of Asp403. Zn2 is coordinated by the carboxyl group of Asp343, Asp323, Lys318 and Glu405. The sulfate ion is within binding proximity to the two zinc ions. **(C)** Close up of N-terminal region where the flexible loop (residues 52-78) of chain X was excised (yellow). Residues flanking excised region 51 and 79 depicted in yellow. The loop is adjacent to the substrate access channel and postulated substrate regulation loop of chain A (red). PDB ID: 6WVV.

The *Pv*-M17Δ53-78 and *Pf*A*-*M17 active site structures align closely, including the bound metal ions. However, while *Pf*A*-*M17 crystals required zinc soaking to occupy both sites, *Pv*-M17Δ53-78 crystals did not require any metal ion treatment. The *Pv*-M17Δ53-78 maps showed clear electron density for two zinc ions in each active site at the same positions observed in the *Pf*A*-*M17 structure (Fig 5B, Supp. Fig S4C). Each of the metal ions was coordinated in a tetrahedral fashion by neighboring active site residues and all residue coordination points were conserved with the *Pf*A*-*M17 structure (Fig 5B). *Pf*A-M17, like most M17 aminopeptidases, has a carbonate ion located within the active site (Supp. Fig S4C) (11). In *Pv*-M17Δ53-78, we observed a sulfate ion in the position of the carbonate (Fig 5B). The presence of the sulfate ion in the *Pv*-M17Δ53-78 active site is likely a crystallization artefact (Fig 5B), given the presence of sulfate ions in the crystallization buffer. Several M17 aminopeptidases have also reported the presence of an active site sulfate ion (50,51), although the vast majority of M17 aminopeptidases bind a carbonate ion at this position. Interactions between the sulfate ion and the zinc ion may contribute to stabilizing metal ion binding in the crystal structure, with atomic distances between the sulfate ion and Zn1 and Zn2 at lengths of 3.1 and 3.3 Å respectively (Supp. Fig S4B).

Kinetic characterization of *Pv*-M17Δ53-78, showed that the enzyme had a faster substrate turnover rate, but 3-fold lower substrate affinity when compared to the wild type, suggesting that the *Pv* loop has a role in substrate regulation (Table 1). The *Pv* loop is adjacent to a large channel through which substrates likely gain access to the buried active site (Fig 5C). With excision of the *Pv* loop, the channel entrance is more exposed, allowing substrates to access the active site more readily and resulting in elevated substrate turnover speed. As excision of the *Pv* loop allows substrates to access the active site more readily, it may similarly allow substrates to leave the active site more readily, causing a decrease in substrate affinity. The channel entrance is flanked by a second loop, contributed by a neighboring chain, that has been described as a ‘substrate regulation loop’ (Fig 5C) (11).

We further characterized the *Pv-*M17 structure using cryo-EM, which enabled us to determine the structure of the full-length protein rather than the loop deletion mutant (PDB ID: 7K5K) (Supp. Table S2). Data was collected from samples prepared in the presence of 1 mM Mn^2+^ to encourage formation of active hexamers and resulted in a map with global resolution of 2.8 Å (Supp. Fig S5) that showed *Pv*-M17 in the conserved hexameric conformation (Fig 6A). Map quality and resolution was the highest at the core of the hexamer and reduced towards the periphery of the hexamer. Each of the active sites had density for two metal ions, which were modelled as Mn^2+^ ions, as well as a single carbonate ion. The carbonate ion electron density was planar in comparison to the *Pv*-M17Δ53-78 sulfate ion tetrahedral electron density observed at the same position (Fig 6B). The position of residues involved in metal ion coordination were conserved between the full length cryo-EM structure and the *Pv*-M17Δ53-78 crystal structure despite the difference in metal ion identity and inclusion of the carbonate ion. The Mn^2+^ ions in the cryo-EM structure were slightly displaced when compared to the Zn^2+^ ions in the crystal structure, although this may be due to the larger atomic radius of Mn^2+^ in comparison to Zn^2+^. The carbonate ion bound in cryo-EM structure active site was positioned 4.3 and 3.7 Å away from Mn1 and Mn2 respectively, distances that are likely beyond the physically feasible Mn-O bond length (Supp. Fig S4D). This indicates that the occupation of both metal ion sites is not reliant on stabilization from the large sulfate ion as observed in the crystal structure.

**Figure 5.**
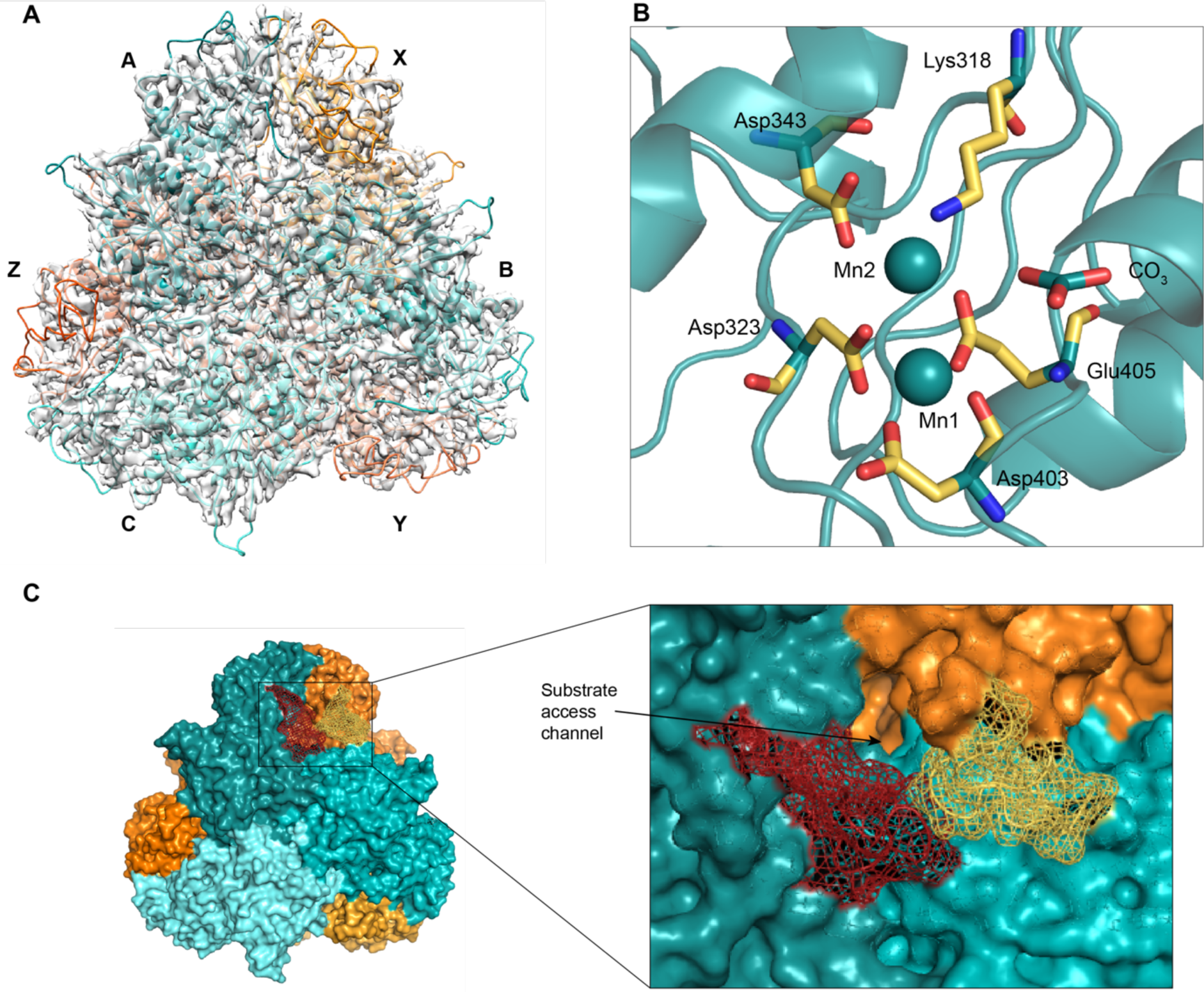
The full length *Pv*-M17 cryo-EM structure also adopts the conserved M17 hexameric conformation and has a conserved active site conformation. **(A**) 2.6 Å cryo-EM structure of full length *Pv-*M17. Cryo-EM structure also adopts the hexameric conformation, consisting of a front trimer depicted in shades of teal and labeled A, B and C, and a back trimer depicted in shades of orange and labeled X, Y and Z. **(B)** *Pv-*M17 full length cryo-EM structure active site. Metal-coordinating residues are shown as yellow sticks, colored by atom. Mn1 and Mn2 are depicted as spheres. Mn1 is coordinated by the carboxyl group of Asp323, Asp403 and Glu405 and the amine group of Asp403. Mn2 is coordinated by the carboxyl group of Asp343, Asp323, Lys318 and Glu405. The active site additional bind a carbonate ion that is beyond binding proximity to the two zinc ions. **(C)** Close up of N-terminal region with *Pv* loop of chain X (yellow) interacting with the substrate regulation loop of chain A (red) and occluding the active site substrate access channel.PDB ID: 7K5K.

While electron density for the *Pv* loop was poorer than the remainder of the hexamer, it could be modelled extending from the exterior of the hexamer into surrounding solvent. In its modelled position, the *Pv* loop is coiled and partially obstructs the substrate access channel (Fig 6C). In solution the *Pv* loop likely extends to occlude more of the channel and further restrict substrate access to the active site, accounting for the disparity in substrate turnover rates between wild type *Pv-*M17 and Pv-M17Δ53-78. The *Pv* loop interacts closely with the substrate regulation loop from the neighboring chain and together the two loops appear to have capacity to influence substrate entry and product egress from the active site. These two loops may function as substrate gatekeepers to the buried active site and operate in concert to regulate both substrate affinity and turnover speed (Fig 6C).

### Structural characterization of the small Pv-M17 oligomers

To gain further insight into the metal-dependent oligomerization mechanism and pathway, we sought to also structurally investigate the small oligomers of *Pv*-M17 that self-associate to form the active hexamer that has been resolved as part of this study (Fig 5A). For this we used cryo-EM, where we could control the environmental metal ion conditions and therefore the dynamic equilibrium and oligomeric species present.

In the presence of 100 mM EDTA, *Pv-*M17 formed a single species corresponding to a tetramer when analyzed using SEC (Supp. Fig S6). We used cryo-EM to investigate the structure of this tetramer. Despite the apparent homogeneity observed on SEC, the EDTA-treated samples were incredibly difficult to optimize for freezing conditions that resulted in good particle distribution across the grid. In total, 56,679 particles were picked from grids, and 16,127 were used to generate a density map with a global resolution of 8.8 Å (Fig 7A, Supp. Fig S5). Molecular dynamic flexible fitting (MDFF) using various combinations of tetramer derived from the hexamer atomic model were used to produce the final model of *Pv*-M17 in the absence of metal (Fig 7A). After MDFF there were some regions of the map where the model fit poorly, which may be a result of the low map resolution or unanticipated restructuring of the tetramer subunits and interactions. The best fit tetramer consisted of a dimer from each side of the hexamer, in a ‘dimer of dimers’ conformation (Fig 7A; A and C from front trimer; X and Y from back trimer). The two dimers interact via the C-terminal at the base of the structure and are in an open conformation at the top (Fig 7B). Although the map resolution was not sufficient to achieve an accurate atomic model, the backbone indicated that the C-terminal domains likely interact through an aromatic tetrad consisting of Trp469 and Try470 with the four Trp residues forming two T-shaped pi-pi interactions (Fig 7B). The *Pv* loop (Pro54 – Ala79) from chains C and Y extend to interact with the neighboring chain, with the *Pv* loop from chain C interacting with chain Y and vice versa. (Fig 7C). Chains C and Y interact with chains A and X through a secondary C-terminal interface that is in close proximity to the active sites of each chain (Fig 7D). A disordered loop from chains C and Y (Ser324 – Met340) extends into the C-terminal domain of chains A and X respectively and comes into close proximity with the active sites of A and X. Simultaneously, the loop pulls Asp323 and Asp343 away from the active sites of C and Y suggesting that the active sites are likely in a non-functional conformation.

**Figure 7.**
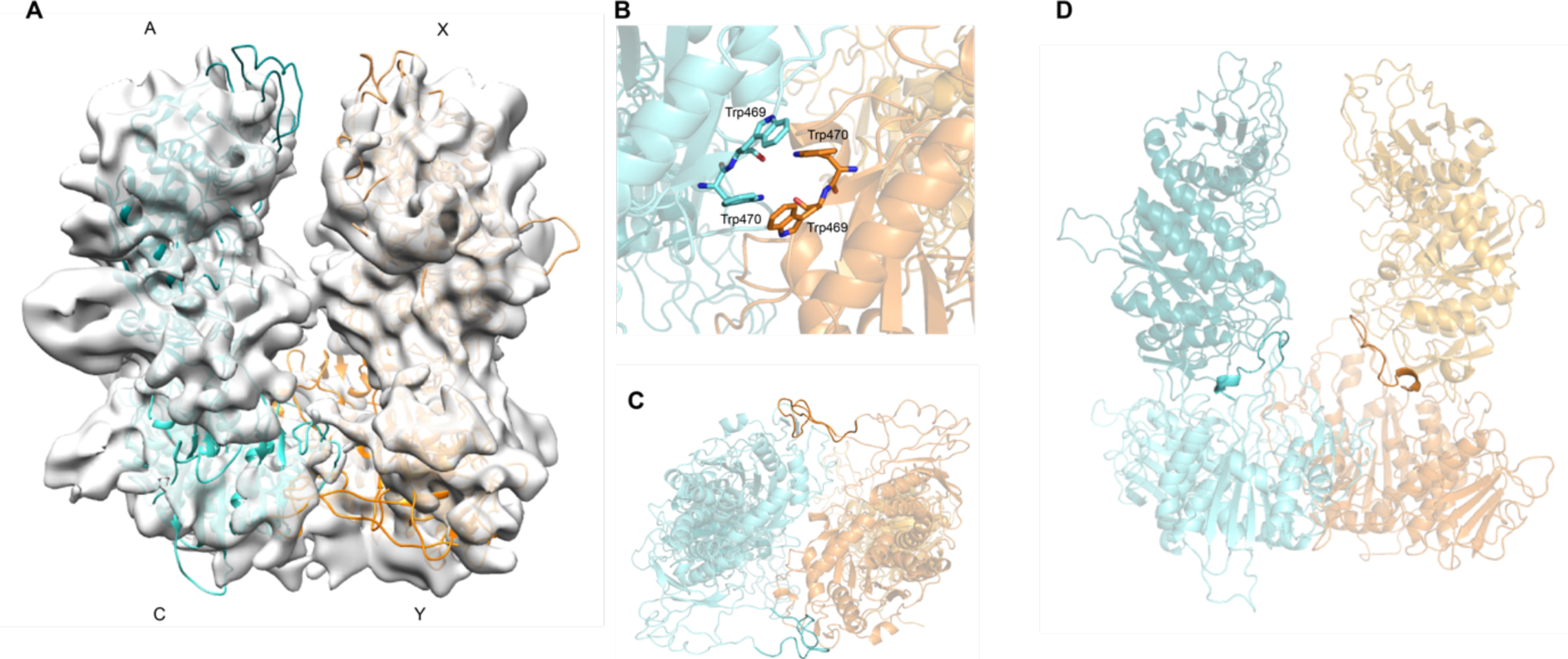
Cryo-EM structure of the *Pv*-M17 tetrameric conformation. **(A)** Tetrameric conformation of *Pv*-M17 adopted in the presence of 100 mM EDTA. Tetramer model was fit to calculated map using molecular dynamic flexible fitting (MDFF). The tetramer adopts a ‘dimer of dimers’ conformation that associate via the C-terminal of chains C and Y. The teal dimer is composed of chains A and C from the ‘front trimer’ of the hexameric conformation, and the orange dimer is composed of chains X and Y from the ‘back trimer’ of the hexameric conformation. **(B)** Proposed aromatic tetrad mediating the C-terminal interface between dimers. Tetrad consists of Trp469 and Trp470 from chain C and Y, each residue forming T-shaped interactions with two others. **(C)** Bottom view of tetramer; the flexible N-terminal loop (residues 53-78) of chains C and Y extend to associate with the neighboring chain and may help to stabilize the C-terminal interface. **(D)** Short C-terminal loops from chains C and Y (residues 324 – 342) extend into chains A and X, coming into close contact with the active sites of chains A and X.

## Discussion

Self-assembly into high order oligomeric conformations is a common mechanism for controlling protein function (52). This form of regulation is involved in countless biological systems; hemoglobin must form α2β2 tetramers to transport oxygen; actin forms long oligomers to facilitate cellular movements (53); porphobilinogen synthase (PBGS) must form homo-octamers to facilitate production of building blocks required to form chlorophyll and vitamin B12 (54). More specifically, many metalloenzymes show a propensity to self-associate into large oligomers; M42 (TET aminopeptidase) (55-57) and M18 (aspartyl aminopeptidase) family aminopeptidases both form tetrahedral dodecamers, while M12 (carboxyl peptidase) family aminopeptidases form tetramers (58). The M17 aminopeptidases are one such family that undergo oligomerization to form homo-hexamers, a process that is essential for catalytic activation of the enzyme.

### Metal-dependent oligomerization: a multi-tiered control mechanism for catalytic activity

Our results indicate that the M17 aminopeptidase oligomerization pathway and formation of the active hexamer is mediated by the acquisition and binding of metal ions within the active site. A similar metal-mediated oligomerization mechanism has previously been described to control the formation of catalytically active M42 dodecamers and PBGS octamers (54-57). Further to controlling the process of M17 hexamerization, both the identity and concentration of metal ions available dictate the extent to which oligomerization occurs. Mn^2+^ and Co^2+^ induced oligomerization of *Pf*A*-*M17 and *Pv*-M17, while Mg^2+^ had little influence and Zn^2+^ caused irreversible protein aggregation. As formation of the hexamer is intrinsically linked to catalytic activity it was expected that maximal catalytic activity would be observed in the presence of Mn^2+^ and Co^2+^, which we saw for both *Pf*A*-*M17 and *Pv-*M17. Similarly, increasing Mn^2+^ concentrations increased formation of hexamer, accompanied by an increase in catalytic activity rates. That we could disrupt this oligomerization/activity relationship by mutating active site residues (PfA-M17(AL) & *Pv-*M17(AL)) confirmed that active site metal binding is central to the oligomerization process. The metal-dependent control of oligomeric states may serve numerous purposes; to prevent unwanted proteolytic damage to cells, to prevent formation of inactive aggregates, or to dictate the role of M17 within an organism. *Plasmodium* in particular are known to have fluctuations in metal ion availability through the life cycle (59), meaning this equilibrium could be implemented as a biological control mechanism, or a metal-dependent ‘activity switch’.

In addition to the metal-dependent oligomerization mechanism controlling activity via formation of the active hexamer, M17 aminopeptidases have a secondary metal-dependent regulation mechanism. The catalysis of the Cys-Gly dipeptide as part of the glutathione breakdown pathway is a postulated role for several bacterial and plant M17 aminopeptidases and involves formation of the active hexamer to carry out the hydrolysis mechanism (60,61). However, Cys-Gly catalysis by *Pf*A*-*M17 and *Pv-*M17 only occurred in the presence of Mn^2+^, and notably did not occur in the presence of other metal ions known to induce hexamerization (e.g. Co^2+^). Cappiello et al. (6) suggested any cations other than Mn^2+^ would interact with the sulfhydryl ion of the Cys residue in such a way that the dipeptide would adopt an unhydrolyzable pose within the active site. Therefore, for Cys-Gly catalysis to occur, M17 aminopeptidases must adopt the hexameric conformation and bind the dipeptide substrate in a hydrolyzable pose; events that are both reliant on active site metal binding.

### Differences in PfA-M17 and Pv-M17 oligomerization behavior is not dictated by active site structure

Throughout this study, we continually observed unexpected differences in metal-regulated oligomerization and activity behaviors between *Pf*A*-*M17 and *Pv-*M17. When the 2.6 Å *Pv-* M17 crystal structure was solved and the active site compared to that of *Pf*A*-*M17, we found that the active site structure was highly conserved and held no indication that metal ions would bind differently between the two structures. The conserved affinity of *Pf*A-M17 and *Pv-*M17 for Leu-Mec reiterates the high degree of active site conservation and indicates that substrate binding is also conserved between the two homologs (*Pf*A*-M17 K*_m_ = 15.0 µM, *Pv-*M17 *K*_m_ = 20.1 µM). Although *Pv-*M17 and *Pf*A*-*M17 metal ion and substrate binding were conserved, *Pv-*M17 exhibited both a faster substrate turnover rate and a greater capacity to form the active hexameric conformation than *Pf*A*-*M17. These different catalytic and oligomeric behaviors are likely due to structural differences at a location distinct from the active site or eventuate from differences in the oligomerization pathway. While the catalytic domains of *Pf*A-M17 and *Pv-*M17 have 90% sequence identity, the N-terminal domains are more divergent with only 53% sequence identity (Supp. Fig S3). The N-termini domains are largely responsible for the intramolecular interactions that likely help to establish and maintain the hexameric conformation. In particular, the N-terminal interface between chains at the ‘top’ of the hexamer, where the chains from opposite sides of the hexamer interact, is considerably more positively charged in *Pv-*M17 than *Pf*A*-*M17. This is largely due to the string of positively charged residues His-Lys-Lys (residues 158-160) in place of the *Pf*A*-*M17 Leu-Ser-Lys. The positively charged regions from each of the N-terminally associating chains sandwich a negatively charged region that, together, contribute to stabilizing this N-terminal interaction. This unique *Pv-*M17 feature is likely just one of many subtle structural differences between the *Pf*A*-*M17 and *Pv-*M17 that culminate in the differences we observe in oligomerization behavior.

### Metal ions drive the M17 dynamic equilibrium towards large oligomeric states

Metal-dependent oligomerization is a key process in dictating the role and function of M17 aminopeptidases, however the process of how M17 aminopeptidases self-associate from monomer to hexamer had not been investigated prior to this study. Our results suggest that in the presence of low concentrations of metal-ion, monomeric M17 aminopeptidase can self-associate to form a dimer, via aromatic interactions at the C-terminal domain interface (Fig 8). Interruption of these aromatic interactions results in the M17 enzyme being locked into a monomeric species, regardless of the metal ion environment. As the environmental metal ion concentration increases, formation of the tetrameric species is observed. This is achieved through association of two free monomers with the existing dimer via C-terminal interactions at a secondary C-terminal interface different to that involved in the dimerization process. Finally, in the presence of sufficient cation, the final two monomers join to form the active hexamer (Fig 8). Based on the MDFF simulated model, the tetramer must undergo slight structural rearrangements to accommodate these final two monomers, which interact with the tetramer species via both C-terminal and N-terminal interactions. This final hexamerization process was disrupted when metal ion binding was compromised, as *Pf*A*-*M17 (AL) and *Pv-* M17 (AL) oligomerized to a tetramer or pentamer but failed to form the hexameric species. This suggests that unimpeded metal binding in the active site is essential for tetramers to undergo the structural rearrangements necessary to form the active hexamer.

**Figure 8.**
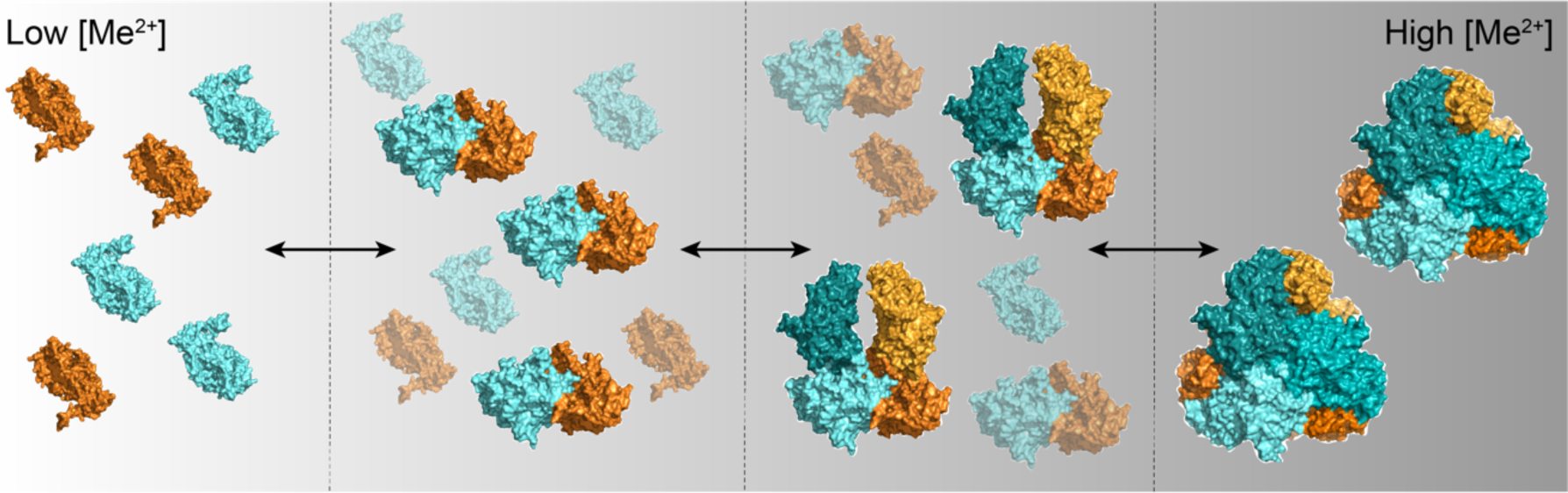
Schematic of the metal-dependent dynamic equilibrium of M17 aminopeptidase oligomeric species. As metal ion concentration increases, the M17 aminopeptidase dynamic equilibrium shifts towards generation of active hexameric species. In low metal ion concentration, monomers self-associate to form dimers. As metal ion concentration increases, monomers associate with the existing oligomers to form tetramers and then hexamers.

We have seen from both AUC and SEC-MALS that *Pv-*M17 and *Pf*A*-*M17 (WT) simultaneously adopt a series of oligomeric species at any one time, indicating that the formation or breakdown of hexamers does not occur via a regimented step-wise fashion with all aminopeptidases adopting the same oligomeric species before progressing to the next. In fact, this would likely be impossible as our oligomerization model suggests the larger oligomeric states are formed by interaction of high order and low order oligomers. Rather, *Pf*A-M17 and *Pv-*M17 exist in a rapid and continuous equilibrium between different oligomeric states that can be controlled by modifying environmental metal ion conditions.

## Conclusions

Beyond their application as antimalarial drug targets (22,23,25), M17 aminopeptidases show potential for use in agricultural (62,63) and environmental industries (64). The ability to adopt numerous oligomeric conformations with distinct functionalities underpins the diversity of potential M17 aminopeptidase applications. The metal-dependent hexamerization mechanism, together with the metal-dependent substrate positioning results in a highly regulated and multi-tiered system with capacity for fine control over catalytic activity. This kind of mechanism lends itself to manipulation of the metal ion environment to dictate enzyme efficiency, substrate specificity and control oligomer-specific functions for pharmaceutical, agricultural and industrial purposes.

## Author Contribution

TRM performed experiments, analyzed data, co-wrote manuscript. SCA and MJB performed experiments, analyzed data. HV and NAB performed experiments, ND performed experiments, analyzed data, concept and SM concept, analyzed data, co-wrote manuscript, provision of funding.

## Acknowledgements

This work was supported by the National Health and Medical Research Council (Synergy Grant 1185354 to SM) and Australian Research Council (DECRA DE190100304 to SA). We thank the Australian Synchrotron (MX-1 & MX-2) and the beamline scientists for beamtime and for technical assistance. We thank the Monash Platforms (Protein Production and Crystallization) for technical assistance. TRM is supported by RTP scholarship from Australian Department of Education and by the Monash Graduate Completion Award.

## Associated content

The following figures and data are provided as supplementary information:

**Supplementary Table S1**. *Pv*-M17 primers used to generate metal-binding mutant and N-terminal loop mutant

**Supplementary Table S2**. Model Parameters of Hexameric *Pv-*M17

**Supplementary Table S3**. Hydrodynamic properties of *Pv-*M17 and *Pf*A*-*M17

**Supplementary Table S4**. *Pv-*M17Δ53-78 X-ray crystal structure data collection and refinement statistics.

**Supplementary Figure S1**. Analytical ultracentrifugation sedimentation equilibrium of *Pf*A-M17 (W525A + Y533A)

**Supplementary Figure S2**. Analytical ultracentrifugation sedimentation equilibrium of *Pv-*M17 in the presence of 50 µM Mn^2+^.

**Supplementary Figure S3**. Sequential alignment with mapped secondary structure of *Pf*A-M17 (PDB ID: 3KQZ) and *Pv*-M17.

**Supplementary Figure S4**. Active site metal ion coordination by *Pv*-M17 and *Pf*A-M17

**Supplementary Figure S5**. Resolution of *Pv-*M17 tetramer and hexamer cryo-EM models

**Supplementary Figure S6**. Size exclusion chromatography trace of *Pv*-M17 forming a tetrameric species in presence of 100 mM EDTA

